# Age-related differences in the impact of background noise on neural speech tracking

**DOI:** 10.1101/2025.01.09.632269

**Authors:** Björn Herrmann

## Abstract

Tracking the envelope of speech in the brain is important for speech comprehension. Recent research suggests that acoustic background noise can enhance neural speech tracking, enabling the auditory system to robustly encode speech even under unfavorable conditions. Aging and hearing loss are associated with internal, neural noise in the auditory system, raising the question whether additional acoustic background noise enhances neural speech tracking in older adults. In the current electroencephalography study, younger (∼25.5 years) and older adults (∼68.5 years) listened to spoken stories in quiet (clear) or in the presence of background noise at a wide range of different signal-to-noise ratios. In younger adults, early neural speech tracking responses (<0.15 s) were enhanced by minimal background noise, indicating response facilitation through noise. In contrast, older adults, compared to younger adults, showed enhanced neural speech tracking for clear speech and speech masked by minimal background noise, but the acoustic noise led to little enhancement of the early neural tracking response in older people. The data demonstrate different sensitivity of the auditory cortex to speech masked by noise between younger and older adults. The results are consistent with the idea that the auditory cortex of older people exhibits more internal, neural noise that enhances neural speech tracking but that additional acoustic noise does not further support speech encoding. The work points to a highly non-linear auditory system that differs between younger and older adults.

## Introduction

Many older adults live with some form of hearing loss (Feder et al., 2015; Goman and Lin, 2016) that leads to difficulties comprehending speech in the presence of background noise, such as in crowded places (Pichora-Fuller et al., 2016; Herrmann and Johnsrude, 2020). Understanding how speech in noisy situations is encoded in the brains of older people is critical for developing effective treatments for speech comprehension challenges.

Much research has focused on how the auditory cortex tracks the envelope of speech (Lalor and Foxe, 2010; Ding et al., 2014; Ding and Simon, 2014; Brodbeck and Simon, 2020), because accurate envelope encoding is thought to support speech understanding (Rosen, 1992; Shannon et al., 1995; Ding et al., 2014; Vanthornhout et al., 2018; Lesenfants et al., 2019). However, recent works suggest non-linearities in how neural speech tracking is affected by different levels of background noise (Yasmin et al., 2023; Herrmann, 2024; Panela et al., 2024). Early envelope tracking responses (<0.15 s) exhibit an inverted u-shaped profile, where the response is highest for moderate signal-to-noise ratios (SNRs) that are associated with intelligible, but challenging speech, whereas the response decreases for more unfavorable SNRs (poor intelligibility) and more favorable SNRs (high intelligibility; Yasmin et al., 2023; Herrmann, 2024). Attentional effort required to understand speech at moderate SNRs has been suggested to lead to the neural-tracking enhancement (Hauswald et al., 2022; Yasmin et al., 2023; Panela et al., 2024), but recent work demonstrates little impact of attention on the inverted u-shape (Herrmann, 2024). Instead, it was suggested that noise per se increases the envelope tracking response at moderate SNRs, and that tracking only decreases when speech intelligibility significantly declines for unfavorable SNRs (Herrmann, 2024). Stochastic resonance – the response facilitation through noise (McDonnell and Abbott, 2009; McDonnell and Ward, 2011; Krauss et al., 2016) – was proposed as the critical mechanism that leads to the neural tracking enhancement (Herrmann, 2024).

Aging and hearing loss are associated with a loss of neural inhibition and an increase in neural excitation in auditory cortex, resulting from reduced inputs to the neural pathway caused by peripheral damage (Caspary et al., 2008; Ouellet and de Villers-Sidani, 2014; Zhao et al., 2016; Resnik and Polley, 2017; Salvi et al., 2017; Herrmann and Butler, 2021; McClaskey, 2024). A loss of inhibition and increased excitation can manifest as hyperresponsivity to sound (Auerbach et al., 2014; Chambers et al., 2016; Salvi et al., 2017). Consistently, the neural tracking of the speech envelope is enhanced in older compared to younger adults (Presacco et al., 2016b, a; Brodbeck et al., 2018; Decruy et al., 2019; Broderick et al., 2021; Panela et al., 2024), highlighting the impact on the encoding of relevant features of speech. Reduced inhibition and increased excitation also increase spontaneous activity – and thus neural noise – in the absence of sound (Kaltenbach and Afman, 2000; Eggermont and Roberts, 2004; Knipper et al., 2013; Eggermont, 2015; Parthasarathy et al., 2019; Knipper et al., 2020). Neural noise in the auditory system – although difficult to observe directly in humans using non-invasive recording techniques, such as electroencephalography (EEG) – could drive age-related enhancements of neural speech tracking through stochastic resonance (for discussions of the role of stochastic resonance in hearing loss see Krauss et al., 2016; Schilling et al., 2023). For example, some neurons may receive insufficient input to elicit a response when an individual listens to clear speech but may be pushed beyond their firing threshold by acoustically elicited neural noise (e.g., in younger) or intrinsic neural noise (e.g., in older) in the auditory system, which, in turn, could lead to increased neural tracking responses. However, whether acoustic background noise at low-to-moderate SNRs, similar to younger adults (Herrmann, 2024), enhances neural speech tracking in the auditory system of older adults is unknown.

The current study uses EEG to investigate in younger and older adults how neural speech tracking is affected by background noise ranging from very high (i.e., intelligible) to low SNRs (i.e., less intelligible). The study’s aim is to elucidate whether external, acoustic background noise enhances the neural tracking response in older adults or whether changes in the aged auditory system reduce noise-driven response facilitations in neural speech tracking.

## Methods and materials

### Participants

Twenty-six younger adults (median: 25.5 years; range: 18–34 years; 8 male or man, 16 female or woman, 1 transgender man, 1 non-binary) and 26 older adults (median: 68.5 years; range: 57–78 years; 9 male or man, 17 female or woman) participated in the current study. Participants were native English speakers or grew up in English-speaking countries (mostly Canada) and have been speaking English since early childhood (<5 years of age). Participants reported having normal hearing abilities and no neurological disease (one person reported having ADHD, but this did not affect their participation). Participants gave written informed consent prior to the experiment and were compensated for their participation. The study was conducted in accordance with the Declaration of Helsinki, the Canadian Tri-Council Policy Statement on Ethical Conduct for Research Involving Humans (TCPS2-2014), and was approved by the Research Ethics Board of the Rotman Research Institute at Baycrest Academy for Research and Education.

### Acoustic environment and stimulus delivery

Data were gathered in a sound-attenuating booth to reduce external sound interference. Sounds were delivered using Sennheiser HD 25-SP II headphones connected via an RME Fireface 400 audio interface. The experiment was implemented using Psychtoolbox (version 3.0.14) running in MATLAB (MathWorks Inc.) on a Lenovo T480 laptop with Windows 7. Visual stimuli were projected into the booth via a mirrored display. Auditory stimuli were played at approximately 70 dB SPL.

### Hearing assessment

Pure-tone audiometry was administered for each participant at frequencies of 0.25, 0.5, 1, 2, 3, 4, 6, and 8 kHz. Pure-tone average thresholds (PTA: average across 0.5, 1, 2, and 4 kHz; Stevens et al., 2013; Humes, 2019) were higher for older compared to younger adults (t_50_ = 6.893, p = 8.8 .10^−9^, d = 1.912; Figure 1A). Elevated thresholds are consistent with the presence of mild-to-moderate hearing loss in the current sample of older adults, as would be expected (Moore, 2007; Plack, 2014; Presacco et al., 2016b; Herrmann et al., 2018, 2022). A few older adults of the current sample also appeared to have ‘clinical’ hearing loss as indicated by thresholds above 20 dB HL (Stevens et al., 2013; Humes, 2019), but none of them were prescribed with hearing aids. Although the main analyses focus on the originally intended comparisons of younger and older adults, in explorative analyses, data were analyzed separately for older adults with clinically ‘normal’ hearing according to standard recommendations (PTA < 20 dB HL; Stevens et al., 2013; Humes, 2019; WHO, 2024) and those with hearing impairment (PTA > 20 dB HL).

**Figure 1:**
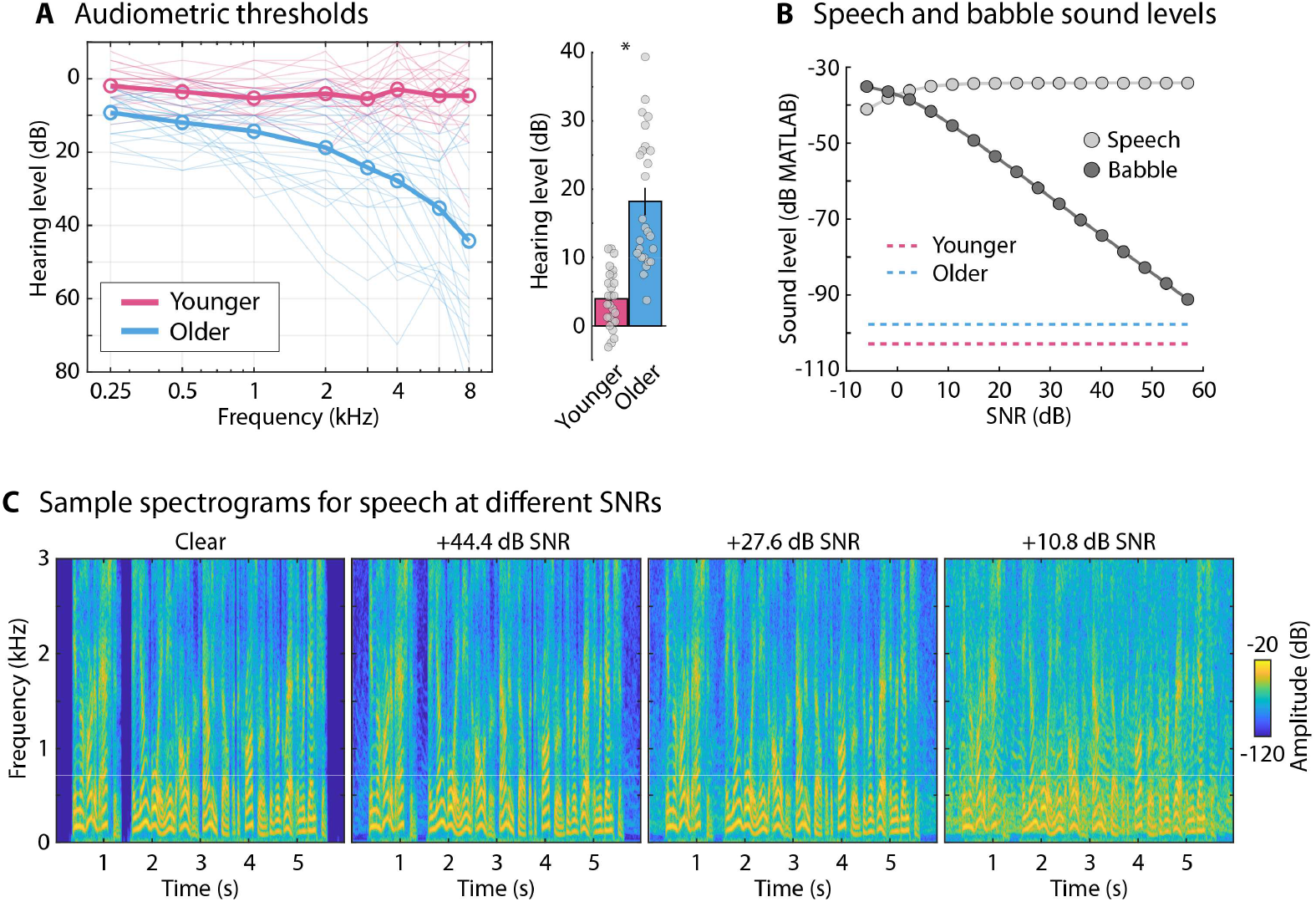
Audiograms, stimulus sound levels, and sample spectrograms. **A:** Left: Pure-tone audiometric thresholds for younger and older adults. Circles and thick lines reflect the mean across participants. Thin lines are the thresholds for each participant. Right: Pure-tone average threshold (PTA; across 0.5, 1, 2, and 4 kHz). Bars reflect the mean across participants. Error bars reflect the standard error of the mean. Dots reflect the PTAs for individual participants. **B:** Sound levels of the speech and background babble for each signal-to-noise ratio (SNR) used in the current study. Sound levels are provided in dB based on MATLAB calculations. More negative values reflect softer sound intensities. Values can be interpreted relative to each other, whereas the absolute magnitude is related to hardware and software conditions, such as sound card, transducers, and MATLAB internal settings. The colored dashed lines show the mean sensation level for a babble noise stimulus for both age groups. **C:** Sample spectrograms for the first 6 seconds of one story under different speech-clarity conditions (clear, 44.4, 27.6, and 10.8 dB SNR). Note that the magnitudes in panels B and C are not comparable.

In order to obtain a reference threshold in MATLAB software for speech and babble presentation during the main experimental procedures, the sensation level for a 12-talker babble noise was estimated using a method-of-limits procedure (Leek, 2011; Herrmann and Johnsrude, 2018; Herrmann et al., 2022). Participants listened to a 14-s babble noise that changed continuously in intensity at a rate of 5.4 dB/s (either decreased [i.e., starting at suprathreshold levels] or increased [i.e., starting at subthreshold levels]). Participants pressed a button when they could no longer hear the noise (intensity decrease) or when they started to hear the tone (intensity increase). The sound stopped after the button press. The sound intensity at the time of the button press was noted for 6 decreasing sounds and 6 increasing sounds (decreasing and increasing sounds alternated), and these were averaged to determine the sensation level. Due to technical issues this threshold was only available for 21 younger and 25 older adults. As expected, given the audiometric pure-tone average thresholds (Figure 1A), sensation levels for the babble noise were elevated for older compared to younger adults (t_44_ = 2.573, p = 0.014, d = 0.762, mean difference: 5.2 dB; Figure 1B).

### Story materials

Participants listened to 20 unique audio stories, each with a duration between 1.5 to 2.5 minutes. These stories were crafted using OpenAI’s GPT-3.5 (OpenAI et al., 2023), which also generated four comprehension questions per story, alongside four answer options (one correct, three distractors). The themes varied widely across stories, encompassing scenarios such as making an unexpected friendship on a plane, a boy finding a knitting talent, and a linguist deciphering ancient text. To ensure high quality of both the content and questions, the AI-generated materials underwent manual verification. Google’s AI-based text-to-speech synthesizer was employed to produce the auditory version of the stories, using the male English-speaking voice “en-US-Neural2-J” with default settings (https://cloud.google.com/text-to-speech/docs/voices).

Participants listened to the 20 stories in 5 blocks, each comprising 4 stories. Four of the 20 stories were played under clear conditions, that is, in quiet. Twelve-talker babble was added to the other 16 stories (Bilger, 1984; Bilger et al., 1984; Wilson et al., 2012). Twelve-talker babble simulates a crowded restaurant, but does not permit identifying individual words in the masker (Mattys et al., 2012). The babble masker was added at SNRs ranging from +57 to –6 dB in 16 steps of 4.2 dB SNR (Figure 1B, C). Speech in background babble above +15 dB SNR is highly intelligible (Holder et al., 2018; Rowland et al., 2018; Spyridakou et al., 2020; Irsik et al., 2022), and listeners had no trouble understanding speech at the highest SNRs used in the current study. All stimuli (clear speech; the mixed speech and babble signals) were normalized to the same root-mean-square amplitude and presented at about 70 dB SPL. Figure 1B shows that for high SNRs, the root-mean-square amplitude of the speech signal in the sound mixture remained relatively constant, because the babble at high SNRs had little impact on the root-mean-square amplitude of the sound mixture. Assignment of speech-clarity levels (clear speech and SNRs) to specific stories was randomized for each participant.

After each story, participants rated two statements regarding their speech comprehension using a 9-point scale (1 = strongly disagree, 9 = strongly agree): ‘I understood the gist of the story’ and ‘I was able to comprehend the speech well’. They were instructed to rate the statements independently from other stories they had heard. Ratings were linearly normalized to a 0 to 1 scale for statistical purposes, making them comparable to proportion-correct measures (Herrmann, 2024; Mathiesen et al., 2024; Panela et al., 2024). Such gist ratings have previously been shown to strongly correlate with word-report speech intelligibility measures (Davis and Johnsrude, 2003; Ritz et al., 2022). The ratings of the two statements were averaged to obtain one comprehension rating per story and participant. After rating the two statements, participants answered four multiple-choice questions about the content of the story. The comprehension questions offered four answer choices (chance level of 25%). The proportion of correct answers was calculated.

### Electroencephalography (EEG) acquisition and preprocessing

A BioSemi system (BioSemi, Netherlands) was used to record electroencephalographic data from 16 Ag/Ag–Cl electrodes (10-20 system) and two additional electrodes, one positioned on the left and one on the right mastoid. Data were recorded at a 1024 Hz sampling rate and with a 208 Hz online low-pass filter. Reference electrodes were part of the BioSemi CMS-DRL (common mode sense-driven right leg) system for optimal referencing and noise reduction.

Offline processing was performed in MATLAB. A 60-Hz elliptic notch filter was used to reduced power-line noise. EEG signals were re-referenced to the average of the left and right mastoid electrodes, which enhances auditory responses at fronto-central electrodes (Ruhnau et al., 2012; Herrmann, 2024). EEG data were high-pass filtered at 0.7 Hz (length: 2449 samples, Hann window) and low-pass filtered at 22 Hz (length: 211 samples, Kaiser window, β = 4). The data were time-locked to the onset of each story, downsampled to 512 Hz, and subjected to an Independent Component Analysis (ICA) to remove blink and eye movement artifacts (Bell and Sejnowski, 1995; Makeig et al., 1995; Oostenveld et al., 2011). Signal segments showing fluctuations greater than 80 μV within a 0.2-second window in any EEG channel were set to 0 μV to remove artifacts not removed by the ICA (cf. Dmochowski et al., 2012; Dmochowski et al., 2014; Cohen and Parra, 2016; Irsik et al., 2022; Yasmin et al., 2023; Panela et al., 2024). Finally, EEG data were further low-pass filtered at 10 Hz (251 points, Kaiser window, β = 4) because previous work has shown that neural signals in this low-frequency range track the speech envelope (Luo and Poeppel, 2007; Di Liberto et al., 2015; Zuk et al., 2021; Karunathilake et al., 2023; Synigal et al., 2023; Yasmin et al., 2023).

### Calculation of amplitude-onset envelopes

Each story’s audio signal (devoid of background noise) was processed through a basic auditory model, which included 30 cochlear-like auditory filters and cochlear compression by a factor of 0.6 (McDermott and Simoncelli, 2011). The resulting 30 envelopes were averaged and smoothed with a 40-Hz low-pass filter (Butterworth, 4^th^ order). Such a computationally simple peripheral model has been shown to be sufficient, as compared to complex, more realistic models, for envelope-tracking approaches (Biesmans et al., 2017). The amplitude-onset envelope was computed since it elicits strong neural speech tracking (Hertrich et al., 2012) and was used in the previous studies in younger adults that showed noise-related enhancements in neural speech tracking (Yasmin et al., 2023; Herrmann, 2024; Panela et al., 2024). The amplitude-onset envelope was obtained by calculating the first derivative of the averaged amplitude envelope and subsequently setting negative values to zero (Hertrich et al., 2012; Fiedler et al., 2017; Daube et al., 2019; Fiedler et al., 2019; Yasmin et al., 2023; Panela et al., 2024). It was then downsampled to match the EEG data’s temporal resolution and transformed to z-scores (subtraction by the mean and division by the standard deviation).

### EEG analysis: Temporal response function and prediction accuracy

The relationship between EEG signals and auditory stimuli was assessed using a linear temporal response function (TRF) model (Crosse et al., 2016; Crosse et al., 2021). Ridge regression with a regularization parameter of λ = 10 was applied based on previous work (Fiedler et al., 2017; Fiedler et al., 2019; Yasmin et al., 2023; Panela et al., 2024). Pre-selection of λ based on previous work avoids extremely low and high λ on some cross-validation iterations and avoids substantially longer computational time. Pre-selection of λ also avoids issues if limited data per condition are available, as in the current study (Crosse et al., 2021). Please note that using other fixed λ (0.01, 0.1, 1, 100) or a cross-correlation approach all led to qualitative similar results.

For each story, 50 random 25-second segments of the EEG data were extracted and paired with corresponding segments of the amplitude-onset envelope. A leave-one-out cross-validation approach was employed, with one segment reserved for testing and the other non-overlapping segments used to train the TRF model for lags ranging from 0 to 0.4 s. The model’s performance was evaluated by correlating the predicted EEG signals with the actual EEG in the test segment, and this procedure was repeated across all 50 segments to derive the mean prediction accuracy. Overlapping segments were used to increase the amount of data for training given the short duration of the stories (Herrmann, 2024). Critically, speech-clarity levels were randomized across stories and analyses were the same for all conditions. Hence, no impact of overlapping training data on the results is expected (consistent with noise-related enhancements observed previously when longer stories and non-overlapping data were used; Yasmin et al., 2023).

To investigate the neural-tracking response amplitude, TRFs for each training dataset were calculated for lags ranging from -0.15 to 0.5 s. Baseline correction was performed by subtracting the mean signal from -0.15 to 0 seconds from the TRF data at each time point. Analysis concentrated on the fronto-central electrodes (F3, Fz, F4, C3, Cz, C4), which are known to reflect auditory cortical activity (Näätänen and Picton, 1987; Picton et al., 2003; Herrmann et al., 2018; Irsik et al., 2021). Key metrics were the P1-N1 and P2-N1 amplitude differences of the TRF. The P1, N1, and P2 latencies were estimated for each SNR from the averaged time courses across participants (separately for each group). P1, N1, and P2 amplitudes were calculated for each participant and condition as the mean amplitude in the 0.02 s time window centered on the peak latency. The P1-minus-N1 and P2-minus-N1 amplitude differences were calculated. The amplitude of individual TRF components (P1, N1, P2) was not analyzed because the TRF time courses for the clear condition had an overall positive shift (see also Herrmann, 2024; Panela et al., 2024) that could bias analyses more favorably towards response differences which may, however, be harder to interpret. The P1-N1 amplitude is of particular interest in the current study, because this early auditory response has previously shown the response amplification through background noise (Yasmin et al., 2023; Herrmann, 2024; Panela et al., 2024).

### Statistical analyses

Behavioral data (comprehension accuracy, comprehension ratings), TRFs, and EEG prediction accuracy for the four clear stories were averaged. For the stories in babble, a sliding average across SNR levels was calculated for behavioral data, TRFs, and EEG prediction accuracy, such that data for three neighboring SNR levels were averaged to reduce noise in the data.

For the statistical analyses of behavioral data (comprehension accuracy, comprehension ratings), P1-N1 amplitude, P2-N1 amplitude, and EEG prediction accuracy, the clear condition was compared to each SNR level (resulting from the sliding average) using a paired samples t-test. False discovery rate (FDR) was used to account for multiple comparisons (Benjamini and Hochberg, 1995; Genovese et al., 2002). Age groups were compared at each SNR level individually using an independent-samples t-test and FDR thresholding. To investigate the overall effect of background babble and interaction with group, a repeated-measures analysis of variance (rmANOVA) was calculated with the within-participants factor Speech Clarity (clear, babble [averaged across SNR levels]) and Group (younger, older). In addition, analyses also explored neural responses for the older adult group split into those with clinically normal hearing (N = 15; PTA < 20 dB HL) and those with hearing impairment (N = 11; PTA > 20 dB HL).

All statistical analyses were carried out using MATLAB (MathWorks) and JASP software (JASP, 2024; version 0.19.1). Note that for post hoc tests of an rmANOVA, JASP uses the rmANOVA degrees of freedom. The reported degrees of freedom may thus be higher than for direct contrasts had they been calculated independently from the rmANOVA.

## Results

### Older adults show reduced noise-related enhancements of neural speech tracking

For both younger and older adults, story comprehension accuracy decreased for the most difficult SNRs relative to clear speech, but there were no differences between age groups (FDR-thresholded; Figure 2A). Ratings of speech comprehension/gist understanding decreased for SNRs below +10 dB compared to clear speech, for both age groups. Older adults also rated speech comprehension/gist understanding higher than younger adults for SNRs between +6.6 and +40.2 dB, which may be related to the known higher subjective ratings of hearing abilities relative to objective hearing abilities in older compared to younger adults (Helfer et al., 2017; Helfer and Jesse, 2021).

**Figure 2:**
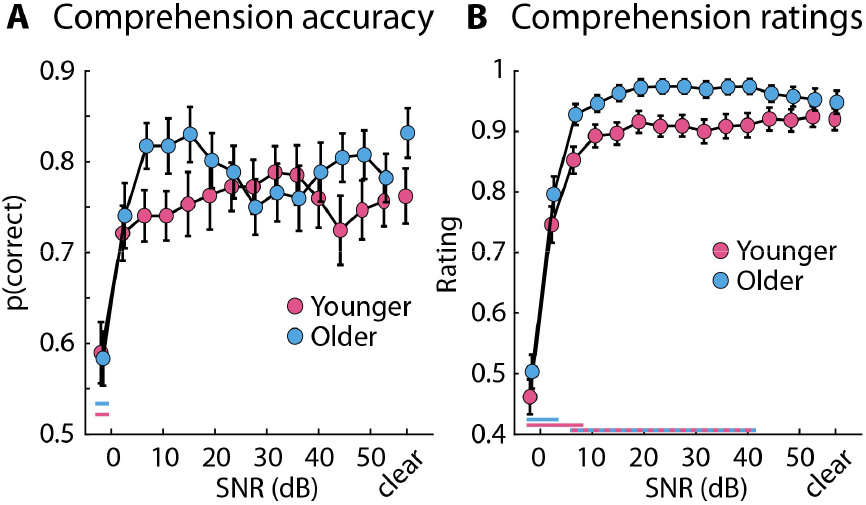
Behavioral results. **A:** Story comprehension accuracy. **B:** Ratings of speech comprehension and gist understanding. The colored, horizontal lines close to the x-axis reflect a significant difference between clear speech and the different SNRs (FDR-thresholded). The two-colored (dashed) line reflects a significant difference between age groups (FDR-thresholded).

Figures 3A and 3B show the temporal responses functions and topographical distributions. Figures 3C displays P1-N1 amplitudes as they relate to speech-clarity conditions. For younger adults, P1-N1 amplitudes increased with decreasing SNR relative to clear speech, up to about +10 dB SNR, whereas amplitudes decreased for yet lower SNRs (Figure 3C, left). For older adults, the increase in P1-N1 amplitudes associated with background babble was only significant around +10 dB SNR, with amplitudes decreasing for lower SNRs (Figure 3C, left). In fact, P1-N1 amplitudes were greater for older compared to younger adults only for clear speech and for speech at high SNRs (i.e., >28 dB), because the auditory cortex of older adults showed a reduced sensitivity to background babble (Figure 3C, left). This reduced noise-sensitivity is also evidenced by the rmANOVA for the P1-N1 amplitude (clear speech vs speech in babble [collapsed across SNRs]). The Speech Clarity × Group interaction (F_1,50_ = 15.714, p = 2.3 · 10^−4^, ω^2^ = 0.025) showed that the P1-N1 amplitude for younger adults was greater for speech in babble than for clear speech (t_25_ = 6.893, p_Holm_ = 2.1 · 10^−5^, d = 0.611), whereas this was not the case for older adults (t_25_ = 0.392, p_Holm_ = 0.697, d = 0.046; Figure 3C, right; effect of Speech Clarity: F_1,50_ = 11.628, p = 0.001, ω^2^ = 0.018; effect of Group: F_1,50_ = 16.207, p = 1.9 · 10^−4^, ω^2^ = 0.13).

**Figure 3:**
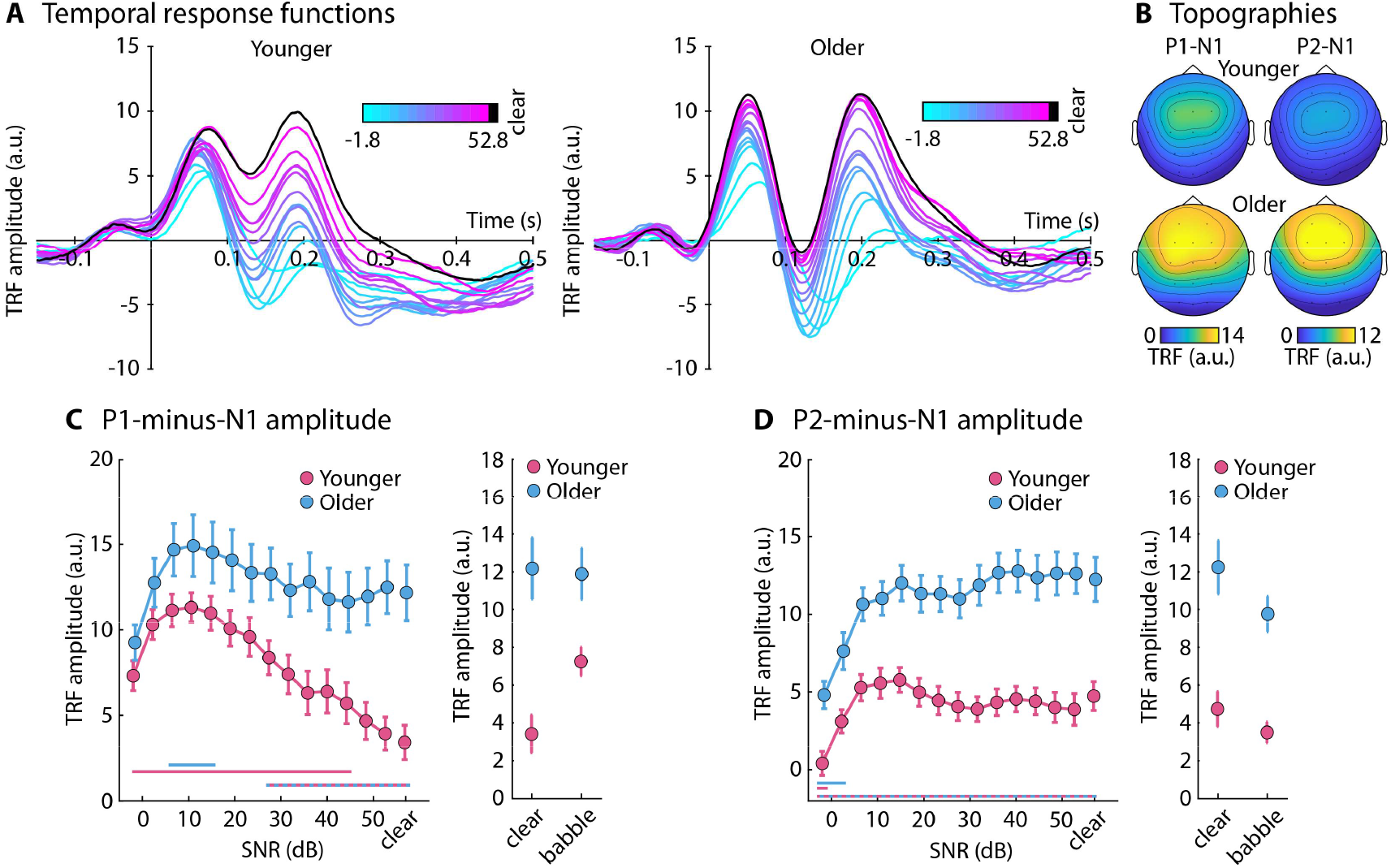
Results for metrics derived from temporal response functions. **A:** Temporal response functions for different speech-clarity conditions and for younger and older adults. **B:** Topographies for P1-N1 and P2-N1 TRF amplitudes. **C:** Left: P1-N1 amplitudes for different speech-clarity conditions and both age groups. The colored, horizontal lines close to the x-axis reflect a significant difference between clear speech and the SNR conditions (FDR-thresholded). The two-colored (dashed) line reflects a significant difference between age groups (FDR-thresholded). Right: P1-N1 TRF amplitude for clear speech and the mean across SNR conditions (babble). **D:** Same as in panel C for the P2-N1 TRF amplitudes.

Figure 3D displays the relation between speech-clarity conditions and P2-N1 amplitudes. For both younger and older adults, the P2-N1 amplitudes were smaller for SNRs around 0 dB and below compared to clear speech, but older adults showed overall larger P2-N1 amplitudes for all speech-clarity conditions. This is also shown by the rmANOVA for the P2-N1 amplitude, revealing smaller amplitudes for speech in babble than clear speech (F_1,50_ = 13.780, p = 5.2 · 10^−4^, ω^2^ = 0.030) and larger amplitudes for older compared to younger adults (F_1,50_ = 26.698, p = 4.2 · 10^−6^, ω^2^ = 0.201). The interaction was not significant (F_1,50_ = 1.540, p = 0.220, ω^2^ = 0.001; Figure 3D).

Figure 4 shows the relation between speech-clarity conditions and EEG prediction accuracy. Prediction accuracy decreased with decreasing SNR relative to clear speech (Figure 4, left). This was also reflected in the rmANOVA, revealing smaller EEG prediction accuracies for speech in babble than clear speech (F_1,50_ = 16.881, p = 1.5 · 10^−4^, ω^2^ = 0.046) and younger compared to older adults (F_1,50_ = 4.832, p = 0.033, ω^2^ = 0.036). The interaction was not significant (F_1,50_ = 0.064, p = 0.801, ω^2^ < 0.001; Figure 4, right).

**Figure 4:**
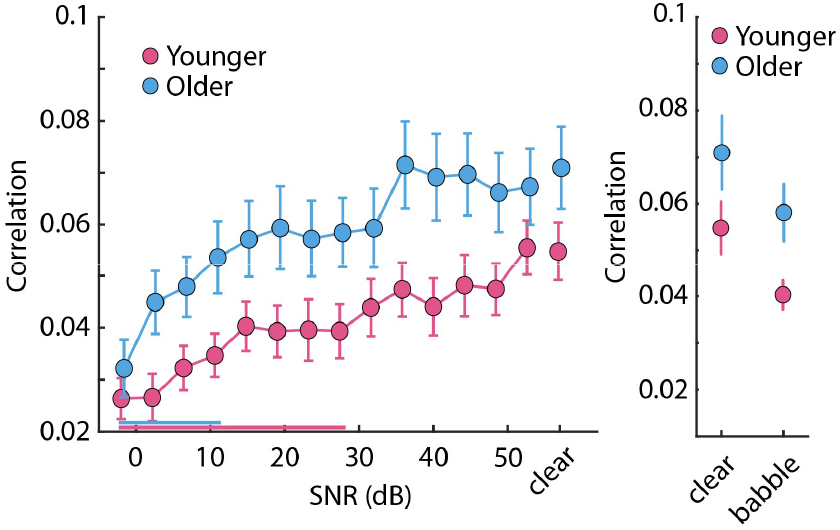
EEG prediction accuracy. **Left:** EEG prediction accuracy for each speech-clarity condition and age group. The colored, horizontal lines close to the x-axis reflect a significant difference between clear speech and the SNR conditions (FDR-thresholded). There was no significant difference between age groups for individual speech-clarity conditions (FDR-thresholded). **Right:** EEG prediction accuracy for clear speech and the mean across SNR conditions (babble).

### Comparing older adults with clinically ‘normal’ hearing to those with hearing impairment

Audiograms for younger adults, older adults with clinically ‘normal’ hearing (definition of PTA < 20 dB HL; WHO, 2024), and older adults with hearing impairment are shown in Figure 5A. Despite the group separation, the older adult group with clinically ‘normal’ hearing still had greater pure-tone average thresholds compared to younger adults (t_39_ = 5.536, p = 2.3 · 10^−6^, d = 1.795) and higher frequency hearing loss, revealing subclinical hearing impairments that are common among older individuals (Dubno et al., 2013; Plack, 2014; Helfer and Jesse, 2021).

**Figure 5:**
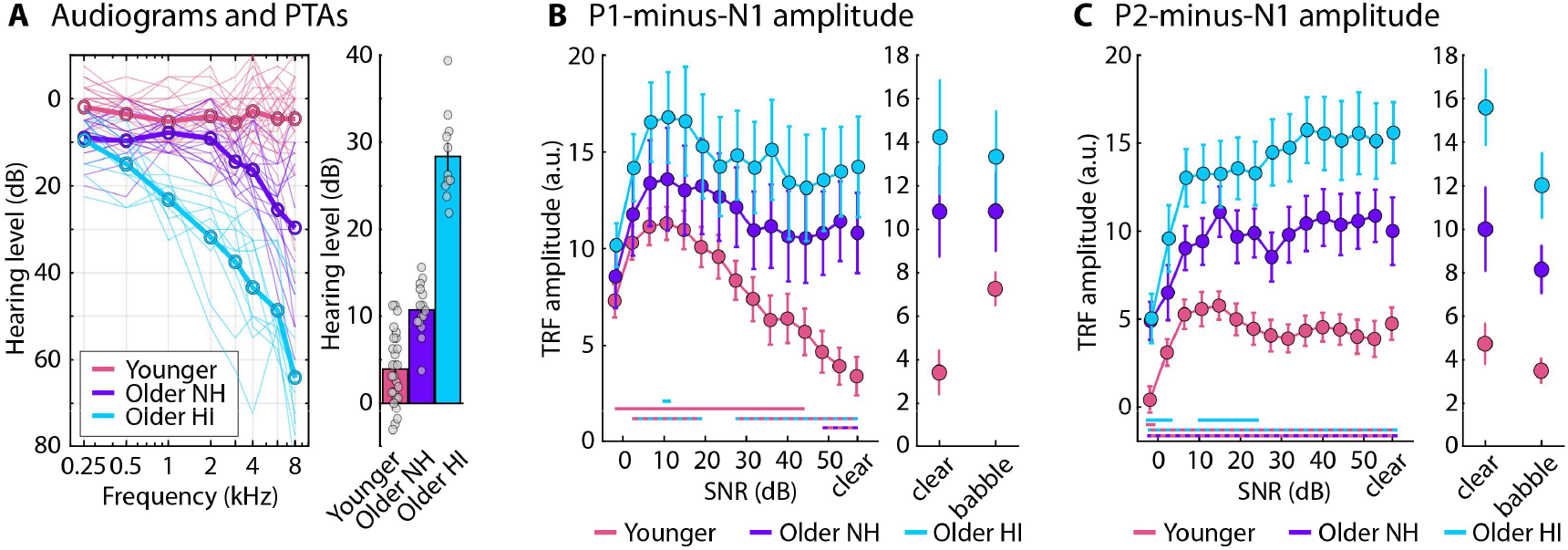
Audiograms and neural-tracking amplitudes for groups split by presence vs absence of hearing impairment. **A:** Audiograms (left) and pure-tone average thresholds for younger adults, older adults with ‘normal’ hearing (NH), and older adults with clinical hearing impairment (HI). **B:** Left: P1-N1 TRF amplitudes for different speech-clarity conditions and groups. The colored, horizontal lines close to the x-axis reflect a significant difference between clear speech and the SNR conditions (FDR-thresholded). The two-colored (dashed) lines reflect a significant difference between groups (FDR-thresholded). Right: P1-N1 TRF amplitude for clear speech and the mean across SNR conditions (babble). **C:** Same as in panel B for the P2-N1 TRF amplitudes.

For neither of the two groups of older adults did the P1-N1 amplitudes show much sensitivity to background babble relative to clear speech, with the exception around +10 dB SNR for older adults with hearing impairment (Figure 5B, left). Critically, for clear speech and speech at high SNRs, the P1-N1 amplitude was larger in both older adult groups compared to younger adults, although the difference was significant for more SNRs for older adults with hearing impairment (FDR-thresholded; Figure 5B). This is also reflected in the rmANOVA for the P1-N1 amplitude, showing a larger amplitude for both groups of older adults compared to younger adults (normal hearing: t_49_ = 2.844; p_Holm_ = 0.013; d = 0.877; hearing impairment: t_49_ = 3.95; p_Holm_ = 7.5 · 10^−4^; d = 1.352), whereas no difference between the two older adult groups was found (t_49_ = 1.256; p_Holm_ = 0.215; d = 0.474; main effect of Group: F_2,49_ = 9.134, p = 4.3 · 10^−4^, ω^2^ = 0.098). Group interacted with Speech Clarity (F_2,49_ = 7.963, p = 0.001, ω^2^ = 0.024; Figure 5B, right): Younger adults showed a larger P1-N1 amplitude when speech was masked by babble compared to clear speech (t_49_ = 5.083; p_Holm_ = 8.7 · 10^−5^; d = 0.614), whereas this was not significant in older adults with normal hearing (t_49_ = 0.009; p_Holm_ = 1; d = 0.001) nor in older adults with hearing impairments (t_49_ = 0.776; p_Holm_ = 1; d = 0.144).

A decrease in P2-N1 amplitudes with SNR, particularly for very low SNRs, relative to clear speech was observed for younger adults and older adults with hearing impairment (FDR-thresholded; Figure 5C left; this was significant for older adults without hearing impairment for uncorrected p-values). Moreover, P2-N1 amplitudes were larger for both older adult groups compared to younger adults for all SNRs (FDR-thresholded; Figure 5C left). The rmANOVA further corroborated this, revealing larger P2-N1 amplitudes for older adults with hearing impairment relative to those with normal hearing (t_49_ = 2.62; p_Holm_ = 0.012; d = 0.967) and younger adults (t_49_ = 5.921; p_Holm_ = 9.3 · 10^−7^; d = 1.98), and larger amplitudes for older adults with normal hearing relative to younger adults (t_49_ = 3.36; p_Holm_ = 0.003; d = 1.013; main effect of Group: F_2,49_ = 18.640, p = 9.5 · 10^−7^, ω^2^ = 0.190). P2-N1 amplitudes were lower for speech in babble than clear speech (F_1,49_ = 17.527, p = 1.2 · 10^−4^, ω^2^ = 0.043), but there was no interaction (F_2,49_ = 1.643, p = 0.204, ω^2^ = 0.003; Figure 5C, right).

For EEG prediction accuracy, an SNR-related decrease relative to clear speech was observed for younger adults and older adults with hearing impairment (FDR-thresholded; Figure 6 left). Prediction accuracy was greater for older adults with hearing impairment relative to younger adults for most speech-clarity conditions (FDR-thresholded; Figure 6 left). The rmANOVA for EEG prediction accuracy showed smaller accuracies for speech in babble than clear speech (F_1,49_ = 15.474, p = 2.6 · 10^−4^, ω^2^ = 0.046). Prediction accuracy was greater for older adults with hearing impairment than younger adults (t_49_ = 3.307; p_Holm_ = 0.005; d = 1.085) and older adults with ‘normal’ hearing (t_49_ = 2.408; p_Holm_ = 0.040; d = 0.872), whereas there was no difference between the two latter groups (t_49_ = 0.721; p_Holm_ = 0.474; d = 0.213; effect of Group: F_2,49_ = 5.546, p = 0.007, ω^2^ = 0.057; Figure 6 right).

**Figure 6:**
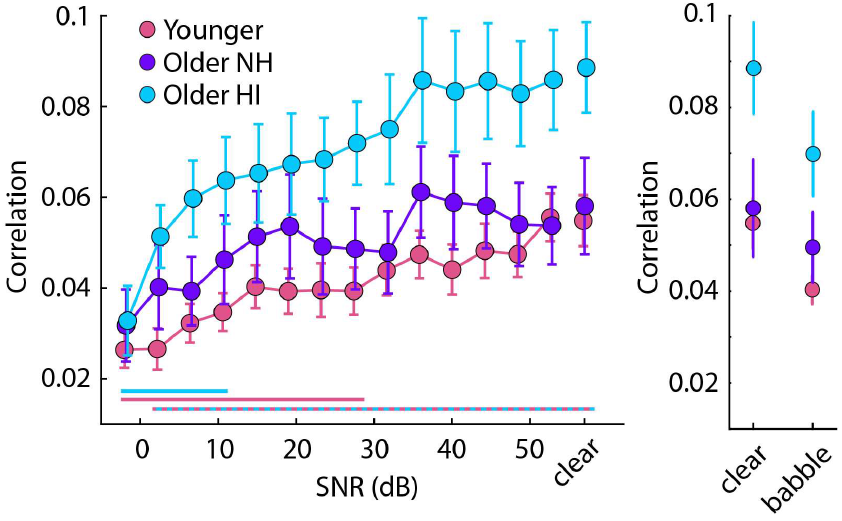
EEG prediction accuracy for groups split by the presence vs absence of hearing impairment. **Left:** EEG prediction accuracy for each speech-clarity condition and age group. The colored, horizontal lines close to the x-axis reflect a significant difference between clear speech and the SNR conditions (FDR-thresholded). The two-colored (dashed) line reflects a significant difference between groups (FDR-thresholded). **Right:** EEG prediction accuracy for clear speech and the mean across SNR conditions (babble). NH – normal hearing; HI – hearing impairment

## Discussion

The current study investigated the extent to which auditory cortex of older adults shows noise-related enhancements in the early neural speech tracking response. Younger and older adults listened to spoken stories either in quiet (clear) or in the presence of background noise. For younger adults, neural speech tracking, as evidenced by the P1-N1 amplitude of the temporal response functions, was enhanced when speech was presented in minimal background noise. Neural tracking of speech in quiet and in minimal background noise was enhanced for older adults compared to younger adults, but older adults showed little evidence of enhancements of the early tracking response due to acoustic background noise. The data indicate a different sensitivity of the auditory cortex between younger and older people to speech masked by acoustic background noise.

### Noise-related enhancement of neural speech tracking

The current study shows that, for younger adults, minimal background noise increases the early response portions of neural tracking of the amplitude-onset envelope of speech compared to speech presented in quiet (P1-N1 amplitude; Figure 3). This is remarkable, given that the background babble overlaps spectrally with the speech, but a noise-related enhancement has been shown recently in a few other works for speech and more simple sounds (Alain et al., 2009; Ward et al., 2010; Parbery-Clark et al., 2011; Alain et al., 2012; Alain et al., 2014; Shukla and Bidelman, 2021; Yasmin et al., 2023; Herrmann, 2024; Panela et al., 2024).

Minor background noise appears to be sufficient to enhance this early neural-tracking response, because the enhancement is present for very high SNRs (>40 dB; Figure 3) for which speech is as intelligible as speech in quiet (Holder et al., 2018; Rowland et al., 2018; Spyridakou et al., 2020; Irsik et al., 2022; Figure 2). Neural speech tracking decreases only for low SNRs for which speech comprehension declines (Figures 2 and 3). Critically, a recent study shows that the noise-related enhancement in speech tracking is present even when participants attend to a demanding visual task rather than to the speech (Herrmann, 2024), making it unlikely that attentional effort explains the enhancement, especially at high SNRs (Rowland et al., 2018). Moreover, the enhancement was present only for early sensory responses (P1-N1), but not for later responses (P2-N1) and EEG prediction accuracy (which integrates responses over time), consistent with a sensory-driven nature of the enhancement.

A few neural speech tracking studies used noise that matched the spectrum of speech as a background masker but did not find a noise-related enhancement (Ding and Simon, 2013; Zou et al., 2019; Synigal et al., 2023). However, these studies used low SNRs for which speech is less intelligible (<10 dB) and the early neural-tracking response is reduced (Figure 3). Moreover, babble noise appears to enhance the neural-tracking response more than speech-matched noise (Herrmann, 2024), potentially because the babble facilitates neural activity in the same speech-relevant auditory regions that are recruited by the speech. This points to some specificity of the spectral noise properties in facilitating the amplification (cf. Krauss and Tziridis, 2021). Other works have used a single competing masker at challenging SNRs (Ding and Simon, 2012; Presacco et al., 2016b, a; Broderick et al., 2018; Kraus et al., 2021; Tune et al., 2021; Teoh et al., 2022), but such maskers fluctuate over time (rather than being stationary as in the current study) and are not very entropic (i.e., noisy).

Stochastic resonance – that is the response facilitation of a non-linear system through noise (Stocks, 2000; Ward et al., 2002; Moss et al., 2004; Stein et al., 2005; McDonnell and Abbott, 2009; McDonnell and Ward, 2011; Krauss et al., 2016; Schilling et al., 2023) – has been suggested to underlie the enhancement of the early neural-tracking response in the presence of background noise (Herrmann, 2024). The term stochastic resonance is increasingly used broadly to describe any phenomenon, not just near-threshold facilitation, where the presence of noise in a nonlinear system improves the quality of the output signal than when noise is absent (McDonnell and Abbott, 2009; McDonnell and Ward, 2011). Speech was presented at suprathreshold levels in the current study, but stochastic resonance may still play a role at the neuronal level (Stocks, 2000; McDonnell and Abbott, 2009). Observing early neural responses to the speech onset-envelope with scalp EEG requires the synchronized activity of more than 10,000 neurons (Niedermeyer and da Silva, 2005; da Silva, 2010). Some neurons may not have been driven enough to elicit a response for speech in quiet and remained near firing threshold, whereas the presence of noise – through stochastic resonance – may have pushed them beyond their firing threshold, thereby explaining the enhancement of the early neural-tracking response (Figure 3C). Stochastic resonance may help individuals to hear robustly even when background noise is present.

### Age-related enhancement in neural speech tracking

Neural speech tracking was enhanced in older compared to younger adults for all metrics (P1-N1, P2-N1, EEG prediction accuracy), especially for speech in quiet (clear) and speech masked by minimal background noise. An age-related enhancement in neural speech tracking is consistent with previous work (Presacco et al., 2016b, a; Brodbeck et al., 2018; Broderick et al., 2021; Panela et al., 2024) and with work showing larger neural responses to tones and noises for older adults (Sörös et al., 2009; Alain et al., 2014; Bidelman et al., 2014; Stothart and Kazanina, 2016; Irsik et al., 2021; Herrmann et al., 2023). Such hyperactivity is thought to result from a loss of inhibition and an increase in excitation in the auditory pathway due to reduced peripheral inputs associated with aging and hearing loss (Caspary et al., 2008; Caspary et al., 2013; Ouellet and de Villers-Sidani, 2014; Zhao et al., 2016; Resnik and Polley, 2017; Salvi et al., 2017; Herrmann and Butler, 2021; McClaskey, 2024).

The age-related enhancement of the P1-N1 amplitude appears mostly to be due to aging (Figure 5B), whereas the P2-N1 amplitude and the EEG prediction accuracy seem to be also or exclusively driven by hearing loss (Figures 5C and 6). Previous studies have shown mixed results regarding the effects of aging versus hearing loss on neural speech tracking. Some studies have found that neural speech tracking is greater for older adults with hearing loss compared to those without (Decruy et al., 2020; Fuglsang et al., 2020; Gillis et al., 2022; Schmitt et al., 2022), whereas other studies point to age-related enhancements per se (Presacco et al., 2019). Counterintuitively, the current data suggest that earlier, sensory responses (P1-N1) are less affected by hearing loss than later responses (P2-N1). However, distinguishing between the impacts of hearing loss versus aging per se may not be possible (Humes et al., 2012). Even minor peripheral damage that is less-well detectable with pure-tone audiometry, such as damage to synapses connecting to auditory nerve fibers (Kujawa and Liberman, 2009; Bharadwaj et al., 2014; Liberman and Kujawa, 2017), can lead to hyperactivity in the auditory system (Qiu et al., 2000; Munguia et al., 2013; Resnik and Polley, 2017; Salvi et al., 2017). The current enhancements for older adults with clinically ‘normal’ hearing compared to younger adults may thus still be related to differences in hearing abilities (i.e., audiometric thresholds were elevated even in normal-hearing older adults; Figure 5A). Speculatively, minor hearing loss is sufficient to enhance early sensory responses, which then do not amplify further with worsening hearing abilities.

### Age-related differences in sensitivity of neural speech tracking to background noise

The main purpose of the current study was to investigate whether auditory cortex of older adults shows enhancements in the early neural-tracking response (P1-N1 amplitude) due to acoustic background noise. However, there was little evidence that the early tracking response in older adults with or without hearing loss is enhanced by acoustic background noise (Figures 3C and 5B). There was only a minor increase in the P1-N1 amplitude around +10 dB SNR, but speech comprehension for this and lower SNRs is more difficult for listeners (Irsik et al., 2022; Herrmann, 2023) and the increase could thus be due to attentional effort (Pichora-Fuller et al., 2016; Hauswald et al., 2022; Yasmin et al., 2023). The early tracking response decreased for both younger and older adults for lower SNRs, for which speech comprehension decreased as well; this is consistent with previous work (Ding and Simon, 2013; Zou et al., 2019; Yasmin et al., 2023).

Story comprehension accuracy did not differ between younger and older adults (Figure 2A), but older adults rated speech comprehension to be easier when speech was minimally (+40 dB SNR) to moderately masked (+10 dB SNR) by background noise, whereas not for speech in quiet or in substantial background noise (Figure 2B). Older adults typically experience more difficulties comprehending speech in noise, although mostly at lower SNRs (Helfer and Freyman, 2008; Ferguson et al., 2010; Presacco et al., 2019; Sobon et al., 2019; Pandey and Herrmann, in press), and the higher ratings for speech in minimal to moderate noise may be due to the known higher subjective ratings of hearing abilities relative to objective hearing abilities in older compared to younger adults (Helfer et al., 2017; Helfer and Jesse, 2021). These differences in behavioral ratings are unlikely to contribute to the age-related reduction in the enhancement of the early tracking response due to background noise. The noise-related response enhancement is unaffected by attention (Herrmann, 2024), and any differences in attentional engagement between age groups is thus unlikely to affect it. Moreover, the fact that the early tracking response for clear speech differed between age groups, whereas behavioral ratings did not, further speaks against a one-to-one mapping of the neural tracking response and behavior.

A loss of neural inhibition and an increase in neural excitation due to aging and hearing loss can lead to a gain or amplification of a neuron’s output, because under such circumstances a smaller input to the neuron is already sufficient for it to fire compared to a less excitable and inhibited neuron. This gain – or central gain, because the amplification increases along the auditory pathway – is often referred to as the cause for hyperresponsivity to sound and speech (Auerbach et al., 2014; Zhao et al., 2016; Salvi et al., 2017; Herrmann and Butler, 2021; McClaskey, 2024). The reduced noise-related enhancement of the early tracking response for older adults could potentially be the result of a maxed-out central gain, such that any noisy neural activity driven by the acoustic noise that would normally facilitate enhancements due to stochastic resonance (such as in younger adults) may not be as effective under conditions of maxed-out central gain.

Alternatively or in addition, spontaneous activity – that is, neural noise – in the auditory system is known to increase with age and hearing loss (Kaltenbach and Afman, 2000; Eggermont and Roberts, 2004; Munguia et al., 2013; Eggermont, 2015; Parthasarathy et al., 2019). Internal, neural noise in older adults may have a comparable effect on the early neural response to speech in quiet as the external, acoustic noise has in younger adults. Indeed, increased neural noise in hearing loss has been suggested to lead to stochastic resonance phenomena where the neural noise can increase sensitivity to sound (Krauss et al., 2016; Krauss and Tziridis, 2021; Schilling et al., 2023). Speculatively, increased internal, neural noise in older adults could perhaps reduce the amplification by external, acoustic noise. That is, if the neurons, that under normal hearing would be near firing threshold, are pushed beyond firing threshold already by the internal noise, then there may be fewer neurons left to be pushed to firing by external, acoustic noise. It is noteworthy, however, that central gain and increased neural noise are not independent phenomena and distinguishing one from the other with non-invasive recording techniques is challenging (Schilling et al., 2023). The current study can thus not distinguish between the two. Future recordings in animal models are needed to further elucidate the cellular mechanisms underlying the reduction of noise-related enhancements of early neural responses in the aged auditory cortex.

## Acknowledgments

I thank Priya Pandey and Saba Junaid for their help with story generation and data collection. The research was supported by the Canada Research Chair program (CRC-2019-00156) and the Natural Sciences and Engineering Research Council of Canada (Discovery Grant: RGPIN-2021-02602).

## Author contributions

**Björn Herrmann:** Conceptualization, methodology, formal analysis, investigation, data curation, writing - original draft, writing - review and editing, visualization, supervision, project administration, funding acquisition.

## Statements and Declarations

The author has no conflicts or competing interests.

## Data availability

Data are available at https://osf.io/ (upon publication). One younger and one older participant declined to share their data publicly (we employ separate consents for study participation and data sharing in line with Canadian Tri-Council Policies for Ethical Conduct for Research Involving Humans – TCPS 2 from 2022). Their data are thus not made available.

## References

Alain C, McDonald K, Van Roon P (2012) Effects of age and background noise on processing a mistuned harmonic in an otherwise periodic complex sound. Hearing Research 283:126–135.

Alain C, Roye A, Salloum C (2014) Effects of age-related hearing loss and background noise on neuromagnetic activity from auditory cortex. Frontiers in Systems Neuroscience 8:Article 8.

Alain C, Quan J, McDonald K, Van Roon P (2009) Noise-induced increase in human auditory evoked neuromagnetic fields. European Journal of Neuroscience 30:132–142.

Auerbach BD, Rodrigues PV, Salvi RJ (2014) Central gain control in tinnitus and hyperacusis. Frontiers in Neurology 5:Article 206.

Bell AJ, Sejnowski TJ (1995) An information maximization approach to blind separation and blind deconvolution. Neural Computation 7:1129–1159.

Benjamini Y, Hochberg Y (1995) Controlling the false discovery rate: a practical and powerful approach to multiple testing. Journal of the Royal Statistical Society Series B 57:289–300.

Bharadwaj HM, Verhulst S, Shaheen L, Liberman MC, Shinn-Cunningham BG (2014) Cochlear neuropathy and the coding of supra-threshold sound. Frontiers in Systems Neuroscience 8:Article 26.

Bidelman GM, Villafuerte JW, Moreno S, Alain C (2014) Age-related changes in the subcorticalecortical encoding and categorical perception of speech. Neurobiology of Aging 35:2526–2540.

Biesmans W, Das N, Francart T, Bertrand A (2017) Auditory-Inspired Speech Envelope Extraction Methods for Improved EEG-Based Auditory Attention Detection in a Cocktail Party Scenario. IEEE Transactions on Neural Systems and Rehabilitation Engineering 25:402–412.

Bilger RC (1984) Manual for the clinical use of the revised SPIN Test. Champaign, IL, USA: The University of Illinois.

Bilger RC, Nuetzel JM, Rabinowitz WM, Rzeczkowski C (1984) Standardization of a Test of Speech Perception in Noise. Journal of Speech, Language, and Hearing Research 27:32–48.

Brodbeck C, Simon JZ (2020) Continuous speech processing. Current Opinion in Physiology 18:25–31.

Brodbeck C, Presacco A, Anderson S, Simon JZ (2018) Over-representation of speech in older adults originates from early response in higher order auditory cortex. Acta Acust United Acust 104:774–777.

Broderick MP, Anderson AJ, Di Liberto GM, Crosse MJ, Lalor EC (2018) Electrophysiological Correlates of Semantic Dissimilarity Reflect the Comprehension of Natural, Narrative Speech. Current Biology 28:803–809.

Broderick MP, Di Liberto GM, Anderson AJ, Rofes A, Lalor EC (2021) Dissociable electrophysiological measures of natural language processing reveal differences in speech comprehension strategy in healthy ageing. Scientific Reports 11:4963.

Caspary DM, Hughes LF, Ling LL (2013) Age-related GABAA receptor changes in rat auditory cortex. Neurobiology of Aging 34:1486–1496.

Caspary DM, Ling L, Turner JG, Hughes LF (2008) Inhibitory neurotransmission, plasticity and aging in the mammalian central auditory system. The Journal of Experimental Biology 211:1781–1791.

Chambers AR, Resnik J, Yuan Y, Whitton JP, Edge AS, Liberman MC, Polley DB (2016) Central Gain Restores Auditory Processing following Near-Complete Cochlear Denervation. Neuron 89:867–879.

Cohen SS, Parra LC (2016) Memorable Audiovisual Narratives Synchronize Sensory and Supramodal Neural Responses. eNeuro 3:e0203.

Crosse MJ, Di Liberto GM, Bednar A, Lalor EC (2016) The Multivariate Temporal Response Function (mTRF) Toolbox: A MATLAB Toolbox for Relating Neural Signals to Continuous Stimuli. Frontiers in human neuroscience 10:604.

Crosse MJ, Zuk NJ, Di Liberto GM, Nidiffer AR, Molholm S, Lalor EC (2021) Linear Modeling of Neurophysiological Responses to Speech and Other Continuous Stimuli: Methodological Considerations for Applied Research. Frontiers in Neuroscience 15.

da Silva FL (2010) EEG: Origin and Measurement. In: EEG - fMRI: Physiological Basis, Technique, and Applications (Mulert C, Lemieux L, eds), pp 19–38. Berlin, Heidelberg: Springer Berlin Heidelberg.

Daube C, Ince RAA, Gross J (2019) Simple Acoustic Features Can Explain Phoneme-Based Predictions of Cortical Responses to Speech. Current Biology 29:1924–1937.

Davis MH, Johnsrude IS (2003) Hierarchical Processing in Spoken Language Comprehension. The Journal of Neuroscience 23:3423–3431.

Decruy L, Vanthornhout J, Francart T (2019) Evidence for enhanced neural tracking of the speech envelope underlying age-related speech-in-noise difficulties. Journal of Neurophysiology 122:601–615.

Decruy L, Vanthornhout J, Francart T (2020) Hearing impairment is associated with enhanced neural tracking of the speech envelope. Hearing Research 393:107961.

Di Liberto Giovanni M, O’Sullivan James A, Lalor Edmund C (2015) Low-Frequency Cortical Entrainment to Speech Reflects Phoneme-Level Processing. Current Biology 25:2457–2465.

Ding N, Simon JZ (2012) Emergence of neural encoding of auditory objects while listening to competing speakers. Proceedings of the National Academy of Sciences 109:11854–11859.

Ding N, Simon JZ (2013) Adaptive temporal encoding leads to a background-insensitive cortical representation of speech. The Journal of Neuroscience 33:5728–5735.

Ding N, Simon JZ (2014) Cortical entrainment to continuous speech: functional roles and interpretations. Frontiers in Human Neuroscience 8.

Ding N, Chatterjee M, Simon JZ (2014) Robust cortical entrainment to the speech envelope relies on the spectro-temporal fine structure. NeuroImage 88:41–46.

Dmochowski JP, Sajda P, Dias J, Parra LC (2012) Correlated components of ongoing EEG point to emotionally laden attention – a possible marker of engagement? Frontiers in Human Neuroscience 6:Article 112.

Dmochowski JP, Bezdek MA, Abelson BP, Johnson JS, Schumacher EH, Parra LC (2014) Audience preferences are predicted by temporal reliability of neural processing. Nature Communications 29:4567.

Dubno JR, Eckert MA, Lee F-S, Matthews LJ, Schmiedt RA (2013) Classifying Human Audiometric Phenotypes of Age-Related Hearing Loss from Animal Models. Journal of the Association for Research in Otolaryngology 14:687–701.

Eggermont JJ (2015) Animal models of spontaneous activity in the healthy and impaired auditory system. Frontiers in Neural Circuits 9:19.

Eggermont JJ, Roberts LE (2004) The neuroscience of tinnitus. Trends in Neurosciences 27:676–682.

Feder K, Michaud D, Ramage-Morin P, McNamee J, Beauregard Y (2015) Prevalence of hearing loss among Canadians aged 20 to 79: Audiometric results from the 2012/2013 Canadian Health Measures Survey. Health Reports 26:18–25.

Ferguson SH, Jongman A, Sereno JA, Keum KA (2010) Intelligibility of Foreign-Accented Speech for Older Adults with and without Hearing Loss. Journal of the American Academy of Audiology 21:153–162.

Fiedler L, Wöstmann M, Herbst SK, Obleser J (2019) Late cortical tracking of ignored speech facilitates neural selectivity in acoustically challenging conditions. Neuroimage 186:33–42.

Fiedler L, Wöstmann M, Graversen C, Brandmeyer A, Lunner T, Obleser J (2017) Single-channel in-ear-EEG detects the focus of auditory attention to concurrent tone streams and mixed speech. Journal of Neural Engineering 14:036020.

Fuglsang SA, Märcher-Rørsted J, Dau T, Hjortkjær J (2020) Effects of sensorineural hearing loss on cortical synchronization to competing speech during selective attention. The Journal of Neuroscience:1936–1919.

Genovese CR, Lazar NA, Nichols T (2002) Thresholding of statistical maps in functional neuroimaging using the false discovery rate. NeuroImage 15:870–878.

Gillis M, Decruy L, Vanthornhout J, Francart T (2022) Hearing loss is associated with delayed neural responses to continuous speech. European Journal of Neuroscience 55:1671–1690.

Goman AM, Lin FR (2016) Prevalence of Hearing Loss by Severity in the United States. American Journal of Public Health 106:1820–1822.

Hauswald A, Keitel A, Chen Y-P, Rösch S, Weisz N (2022) Degradation levels of continuous speech affect neural speech tracking and alpha power differently. European Journal of Neuroscience 55:3288–3302.

Helfer KS, Freyman RL (2008) Aging and Speech-on-Speech Masking. Ear & Hearing 29:87–98.

Helfer KS, Jesse A (2021) Hearing and speech processing in midlife. Hearing Research 402:108097.

Helfer KS, Merchant GR, Wasiuk PA (2017) Age-Related Changes in Objective and Subjective Speech Perception in Complex Listening Environments. Journal of Speech, Language, and Hearing Research 60:3009–3018.

Herrmann B (2023) The perception of artificial-intelligence (AI) based synthesized speech in younger and older adults. International Journal of Speech Technology 26:395–415.

Herrmann B (2024) Minimal background noise enhances neural speech tracking: Evidence of stochastic resonance. BioRxiv.

Herrmann B, Johnsrude IS (2018) Attentional State Modulates the Effect of an Irrelevant Stimulus Dimension on Perception. Journal of Experimental Psychology: Human Perception and Performance 44:89–105.

Herrmann B, Johnsrude IS (2020) A Model of Listening Engagement (MoLE). Hearing Research 397:108016.

Herrmann B, Butler BE (2021) Hearing Loss and Brain Plasticity: The Hyperactivity Phenomenon. Brain Structure & Function 226:2019–2039.

Herrmann B, Maess B, Johnsrude IS (2018) Aging Affects Adaptation to Sound-Level Statistics in Human Auditory Cortex. The Journal of Neuroscience 38:1989–1999.

Herrmann B, Maess B, Johnsrude IS (2022) A Neural Signature of Regularity in Sound is Reduced in Older Adults. Neurobiology of Aging 109:1–10.

Herrmann B, Maess B, Johnsrude IS (2023) Sustained responses and neural synchronization to amplitude and frequency modulation in sound change with age. Hearing Research 428:108677.

Hertrich I, Dietrich S, Trouvain J, Moos A, Ackermann H (2012) Magnetic brain activity phase-locked to the envelope, the syllable onsets, and the fundamental frequency of a perceived speech signal. Psychophysiology 49:322–334.

Holder JT, Levin LM, Gifford RH (2018) Speech Recognition in Noise for Adults With Normal Hearing: Age-Normative Performance for AzBio, BKB-SIN, and QuickSIN. Otology & Neurotology 39:e972–e978.

Humes LE (2019) Examining the Validity of the World Health Organization’s Long-Standing Hearing Impairment Grading System for Unaided Communication in Age-Related Hearing Loss. American Journal of Audiology 28:810–818.

Humes LE, Dubno JR, Gordon-Salant S, Lister JJ, Cacace AT, Cruickshanks KJ, Gates GA, Wilson RH, Wingfield A (2012) Central Presbycusis: A Review and Evaluation of the Evidence. Journal of the American Academy of Audiology 23:635–666.

Irsik VC, Johnsrude IS, Herrmann B (2022) Neural activity during story listening is synchronized across individuals despite acoustic masking. Journal of Cognitive Neuroscience 34:933–950.

Irsik VC, Almanaseer A, Johnsrude IS, Herrmann B (2021) Cortical Responses to the Amplitude Envelopes of Sounds Change with Age. The Journal of Neuroscience 41:5045–5055.

JASP (2024) JASP [Computer software]. In: https://jasp-stats.org/.

Kaltenbach JA, Afman CE (2000) Hyperactivity in the dorsal cochlear nucleus after intense sound exposure and its resemblance to tone-evoked activity: a physiological model for tinnitus. Hearing Research 140:165–172.

Karunathilake IMD, Dunlap JL, Perera J, Presacco A, Decruy L, Anderson S, Kuchinsky SE, Simon JZ (2023) Effects of aging on cortical representations of continuous speech. Journal of Neurophysiology 129:1359–1377.

Knipper M, Van Dijk P, Nunes I, Rüttiger L, Zimmermann U (2013) Advances in the neurobiology of hearing disorders: Recent developments regarding the basis of tinnitus and hyperacusis. Progress in Neurobiology 111:17–33.

Knipper M, van Dijk P, Schulze H, Mazurek B, Krauss P, Scheper V, Warnecke A, Schlee W, Schwabe K, Singer W, Braun C, Delano PH, Fallgatter AJ, Ehlis A-C, Searchfield GD, Munk MHJ, Baguley DM, Rüttiger L (2020) The Neural Bases of Tinnitus: Lessons from Deafness and Cochlear Implants. The Journal of Neuroscience 40:7190–7202.

Kraus F, Tune S, Ruhe A, Obleser J, Wöstmann M (2021) Unilateral Acoustic Degradation Delays Attentional Separation of Competing Speech. Trends in Hearing 25:23312165211013242.

Krauss P, Tziridis K (2021) Simulated transient hearing loss improves auditory sensitivity. Scientific Reports 11:14791.

Krauss P, Tziridis K, Metzner C, Schilling A, Hoppe U, Schulze H (2016) Stochastic Resonance Controlled Upregulation of Internal Noise after Hearing Loss as a Putative Cause of Tinnitus-Related Neuronal Hyperactivity. Frontiers in Neuroscience 10:Article 597.

Kujawa SG, Liberman MC (2009) Adding Insult to Injury: Cochlear Nerve Degeneration after “Temporary” Noise-Induced Hearing Loss. The Journal of Neuroscience 29:14077–14085.

Lalor EC, Foxe JJ (2010) Neural responses to uninterrupted natural speech can be extracted with precise temporal resolution. European Journal of Neuroscience 31:189–193.

Leek MR (2011) Adaptive procedures in psychophysical research. Perception & Psychophysics 63:1279–1292.

Lesenfants D, Vanthornhout J, Verschueren E, Decruy L, Francart T (2019) Predicting individual speech intelligibility from the cortical tracking of acoustic- and phonetic-level speech representations. Hearing Research 380:1–9.

Liberman MC, Kujawa SG (2017) Cochlear synaptopathy in acquired sensorineural hearing loss: Manifestations and mechanisms. Hearing Research 349:138–147.

Luo H, Poeppel D (2007) Phase Patterns of Neuronal Responses Reliably Discriminate Speech in Human Auditory Cortex. Neuron 54:1001–1010.

Makeig S, Bell AJ, Jung T-P, Sejnowski TJ (1995) Independent component analysis of electroencephalographic data. In: Advances in Neural Information Processing Systems (Touretzky D, Mozer M, Hasselmo M, eds), pp 145–151. Cambridge, MA, USA: MIT Press.

Mathiesen SL, Van Hedger SC, Irsik VC, Bain MM, Johnsrude IS, Herrmann B (2024) Exploring age differences in absorption and enjoyment during story listening. Psychology International 6:667–684.

Mattys SL, Davis MH, Bradlow AR, Scott SK (2012) Speech recognition in adverse conditions: A review. Language and Cognitive Processes 27:953–978.

McClaskey CM (2024) Neural hyperactivity and altered envelope encoding in the central auditory system: Changes with advanced age and hearing loss. Hearing Research 442:108945.

McDermott Josh H, Simoncelli Eero P (2011) Sound Texture Perception via Statistics of the Auditory Periphery: Evidence from Sound Synthesis. Neuron 71:926–940.

McDonnell MD, Abbott D (2009) What Is Stochastic Resonance? Definitions, Misconceptions, Debates, and Its Relevance to Biology. PLOS Computational Biology 5:e1000348.

McDonnell MD, Ward LM (2011) The benefits of noise in neural systems: bridging theory and experiment. Nature Reviews Neuroscience 12:415–425.

Moore BCJ (2007) Cochlear Hearing Loss: Physiological, Psychological and Technical Issues. West Sussex, Engand: John Wiley & Sons, Ltd.

Moss F, Ward LM, Sannita WG (2004) Stochastic resonance and sensory information processing: a tutorial and review of application. Clinical Neurophysiology 115:267–281.

Munguia R, Pienkowski M, Eggermont JJ (2013) Spontaneous firing rate changes in cat primary auditory cortex following long-term exposure to non-traumatic noise: Tinnitus without hearing loss? Neuroscience Letters 546:46–50.

Näätänen R, Picton TW (1987) The N1 wave of the human electric and magnetic response to sound: a review and an analysis of the component structure. Psychophysiology 24:375–425.

Niedermeyer E, da Silva FHL (2005) Electroencephalography: Basic Principles, Clinical Applications, and Related Fields: Lippincott Williams & Wilkins.

Oostenveld R, Fries P, Maris E, Schoffelen JM (2011) FieldTrip: Open source software for advanced analysis of MEG, EEG, and invasive electrophysiological data. Computational Intelligence and Neuroscience 2011:Article ID 156869.

Ouellet L, de Villers-Sidani E (2014) Trajectory of the main GABAergic interneuron populations from early development to old age in the rat primary auditory cortex. Frontiers in Neuroanatomy 8:Article 40.

Pandey PR, Herrmann B (in press) The influence of semantic context on the intelligibility benefit from speech glimpses in younger and older adults. Journal of Speech, Language, and Hearing Research.

Panela RA, Copelli F, Herrmann B (2024) Reliability and generalizability of neural speech tracking in younger and older adults. Neurobiology of Aging 134:165–180.

Parbery-Clark A, Marmel F, Bair J, Kraus N (2011) What subcortical–cortical relationships tell us about processing speech in noise. European Journal of Neuroscience 33:549–557.

Parthasarathy A, Herrmann B, Bartlett EL (2019) Aging alters envelope representations of speech-like sounds in the inferior colliculus. Neurobiology of Aging 73:30–40.

Pichora-Fuller MK, Kramer SE, Eckert MA, Edwards B, Hornsby BWY, Humes LE, Lemke U, Lunner T, Matthen M, Mackersie CL, Naylor G, Phillips NA, Richter M, Rudner M, Sommers MS, Tremblay KL, Wingfield A (2016) Hearing Impairment and Cognitive Energy: The Framework for Understanding Effortful Listening (FUEL). Ear & Hearing 37 Suppl 1:5S–27S.

Picton TW, John SM, Dimitrijevic A, Purcell DW (2003) Human auditory steady-state responses. International Journal of Audiology 42:177–219.

Plack CJ (2014) The sense of hearing. New York, USA: Psychology Press.

Presacco A, Simon JZ, Anderson S (2016a) Effect of informational content of noise on speech representation in the aging midbrain and cortex. Journal of Neurophysiology 116:2356–2367.

Presacco A, Simon JZ, Anderson S (2016b) Evidence of degraded representation of speech in noise, in the aging midbrain and cortex. Journal of Neurophysiology 116:2346–2355.

Presacco A, Simon JZ, Anderson S (2019) Speech-in-noise representation in the aging midbrain and cortex: Effects of hearing loss. PLoS ONE 14:e0213899.

Qiu C, Salvi R, Ding D, Burkard R (2000) Inner hair cell loss leads to enhanced response amplitudes in auditory cortex of unanesthetized chinchillas: evidence for increased system gain. Hearing research 139:153–171.

Resnik J, Polley DB (2017) Fast-spiking GABA circuit dynamics in the auditory cortex predict recovery of sensory processing following peripheral nerve damage. eLife 6:e21452.

Ritz H, Wild CJ, Johnsrude IS (2022) Parametric cognitive load reveals hidden costs in the neural processing of perfectly intelligible degraded speech. The Journal of Neuroscience 42:4619–4628.

Rosen S (1992) Temporal Information in Speech: Acoustic, Auditory and Linguistic Aspects. Philosophical Transactions: Biological Sciences 336:367–373.

Rowland SC, Hartley DEH, Wiggins IM (2018) Listening in Naturalistic Scenes: What Can Functional Near-Infrared Spectroscopy and Intersubject Correlation Analysis Tell Us About the Underlying Brain Activity? Trends in Hearing 22:2331216518804116.

Ruhnau P, Herrmann B, Schröger E (2012) Finding the right control: The mismatch negativity under investigation. Clinical Neurophysiology 123:507–512.

Salvi R, Sun W, Ding D, Chen G-D, Lobarinas E, Wang J, Radziwon K, Auerbach BD (2017) Inner Hair Cell Loss Disrupts Hearing and Cochlear Function Leading to Sensory Deprivation and Enhanced Central Auditory Gain. Frontiers in Neuroscience 10:Article 621.

Schilling A, Sedley W, Gerum R, Metzner C, Tziridis K, Maier A, Schulze H, Zeng F-G, Friston KJ, Krauss P (2023) Predictive coding and stochastic resonance as fundamental principles of auditory phantom perception. Brain 146:4809–4825.

Schmitt R, Meyer M, Giroud N (2022) Better speech-in-noise comprehension is associated with enhanced neural speech tracking in older adults with hearing impairment. Cortex 151:133–146.

Shannon RV, Zeng F-G, Kamath V, Wygonski J, Ekelid M (1995) Speech Recognition with Primarily Temporal Cues. Science 270:303–304.

Shukla B, Bidelman GM (2021) Enhanced brainstem phase-locking in low-level noise reveals stochastic resonance in the frequency-following response (FFR). Brain Research 1771:147643.

Sobon KA, Taleb NM, Buss E, Grose JH, Calandruccio L (2019) Psychometric function slope for speech-in-noise and speech-in-speech: Effects of development and aging. The Journal of the Acoustical Society of America 145:EL284–EL290.

Sörös P, Treismann IK, Manemann E, Lütkenhöner B (2009) Auditory temporal processing in healthy aging: a magnetoencephalographic study. BMC Neuroscience 10:34.

Spyridakou C, Rosen S, Dritsakis G, Bamiou D-E (2020) Adult normative data for the speech in babble (SiB) test. International Journal of Audiology 59:33–38.

Stein RB, Gossen ER, Jones KE (2005) Neuronal variability: noise or part of the signal? Nature Reviews Neuroscience 6:389–397.

Stevens G, Flaxman S, Brunskill E, Mascarenhas M, Mathers CD, Finucane M, on behalf of the Global Burden of Disease Hearing Loss Expert G (2013) Global and regional hearing impairment prevalence: an analysis of 42 studies in 29 countries. European Journal of Public Health 23:146–152.

Stocks NG (2000) Suprathreshold Stochastic Resonance in Multilevel Threshold Systems. Physical Review Letters 84:2310–2313.

Stothart G, Kazanina N (2016) Auditory perception in the aging brain: the role of inhibition and facilitation in early processing. Neurobiology of Aging 47:23–24.

Synigal SR, Anderson AJ, Lalor EC (2023) Electrophysiological indices of hierarchical speech processing differentially reflect the comprehension of speech in noise. BioRxiv.

Teoh ES, Ahmed F, Lalor EC (2022) Attention Differentially Affects Acoustic and Phonetic Feature Encoding in a Multispeaker Environment. The Journal of Neuroscience 42:682.

Tune S, Alavash M, Fiedler L, Obleser J (2021) Neural attentional-filter mechanisms of listening success in middle-aged and older individuals. Nature Communications 12:4533.

Vanthornhout J, Decruy L, Wouters J, Simon JZ, Francart T (2018) Speech Intelligibility Predicted from Neural Entrainment of the Speech Envelope. Journal of the Association for Research in Otolaryngology 19:181–191.

Ward LM, Neiman A, Moss F (2002) Stochastic resonance in psychophysics and in animal behavior. Biological Cybernetics 87:91–101.

Ward LM, MacLean SE, Kirschner A (2010) Stochastic Resonance Modulates Neural Synchronization within and between Cortical Sources. PLoS ONE 5:e14371.

WHO (2024) Deafness. In. https://www.who.int/news-room/fact-sheets/detail/deafness-and-hearing-loss.

Wilson RH, McArdle RA, Watts KL, Smith SL (2012) The Revised Speech Perception in Noise Test (R-SPIN)in a Multiple Signal-to-Noise Ratio Paradigm. Journal of the American Academy of Audiology 23:590–605.

Yasmin S, Irsik VC, Johnsrude IS, Herrmann B (2023) The effects of speech masking on neural tracking of acoustic and semantic features of natural speech. Neuropsychologia 186:108584.

Zhao Y, Song Q, Li X, Li C (2016) Neural Hyperactivity of the Central Auditory System in Response to Peripheral Damage. Neural Plasticity 2016:2162105.

Zou J, Feng J, Xu T, Jin P, Luo C, Zhang J, Pan X, Chen F, Zheng J, Ding N (2019) Auditory and language contributions to neural encoding of speech features in noisy environments. NeuroImage 192:66–75.

Zuk NJ, Murphy JW, Reilly RB, Lalor EC (2021) Envelope reconstruction of speech and music highlights stronger tracking of speech at low frequencies. PLOS Computational Biology 17:e1009358.

